# 3D-Printed Biodegradable Microswimmer for Drug Delivery and Targeted Cell Labeling

**DOI:** 10.1101/379024

**Authors:** Hakan Ceylan, I. Ceren Yasa, Oncay Yasa, A. Fatih Tabak, Joshua Giltinan, Metin Sitti

## Abstract

Miniaturization of interventional medical devices can leverage minimally invasive technologies by enabling operational resolution at cellular length scales with high precision and repeatability. Untethered micron-scale mobile robots can realize this by navigating and performing in hard-to-reach, confined and delicate inner body sites. However, such a complex task requires an integrated design and engineering strategy, where powering, control, environmental sensing, medical functionality and biodegradability need to be considered altogether. The present study reports a hydrogel-based, biodegradable microrobotic swimmer, which is responsive to the changes in its microenvironment for theranostic cargo delivery and release tasks. We design a double-helical magnetic microswimmer of 20 μm length, which is 3D-printed with complex geometrical and compositional features. At normal physiological concentrations, matrix metalloproteinase-2 (MMP-2) enzyme can entirely degrade the microswimmer body in 118 h to solubilized non-toxic products. The microswimmer can respond to the pathological concentrations of MMP-2 by swelling and thereby accelerating the release kinetics of the drug payload. Anti-ErbB 2 antibody-tagged magnetic nanoparticles released from the degraded microswimmers serve for targeted labeling of SKBR3 breast cancer cells to realize the potential of medical imaging of local tissue sites following the therapeutic intervention. These results represent a leap forward toward clinical medical microrobots that are capable of sensing, responding to the local pathological information, and performing specific therapeutic and diagnostic tasks as orderly executed operations using their smart composite material architectures.

## INTRODUCTION

Advancements in interventional technologies have enabled minimally invasive strategies, *e.g*., endoscopy or robot-assisted surgery, which have markedly reduced centi/decimeter-size incisions of many open surgeries to millimeter-size holes, lowered post-operative patient morbidity, and shortened recovery times (*1, 2*). The advances and evolution of untethered mobile robots, whose overall size is down to around a single cell, can further leverage minimally invasive medicine by providing an unprecedented direct access and precise control in deep and delicate body sites, such as the central nervous system, the circulatory system, the fetus, and the eye (*3–8*). Recent progress along this line has already resulted in a number of synthetic and biohybrid microrobotic designs with intriguing functionalities toward their use in various body sites (*9–12*).

Active navigation of highly concentrated therapeutic or diagnostic agents to the site of action could represent a state-of-the-art application of microscopic robots, compared to the limited delivery and distribution efficiencies of systemic routes and local diffusion (*13, 14*). By active navigation, it is possible to minimize the effects of systemic toxicity (*15*). Active delivery and tunable release modalities would increase the overall efficacy of single dose administration. Coupled sensing -of local cues in the living environment, *e.g*., disease markers- and trigger -of the release of multiple payload types with programmable kinetics- could pave the way for microrobotic therapy and diagnosis in the form of an orderly executed operation (*16*). A conventional robot responds to the changes in its environment by means of its on-board sensors and computational capabilities. Achieving such capabilities at the smaller dimensions, where such computational capabilities do not exist, is a major research question (*17*). Microorganisms, such as slime molds and bacteria have evolved to use physical intelligence as the main route of making decisions in complex and evolving conditions (*18*). Consequently, programmed physical and chemical properties of materials can enable a robust design route for making microrobotic systems with the capabilities of motion, sensing, and functioning in response to local changes in the environment (*19, 20*).

Biodegradability, *i.e*., decomposition over time as a result of the resident biological activity, is a critical aspect of microrobotic design for their safe operation in the living environment. When the prescribed task is accomplished, the safest option for removing the microrobots from the body is their degradation to non-toxic, metabolized products. The use of non-degradable materials can result in serious acute and chronic toxicities, which could require surgical revision, and hence lower the overall desired benefit from the microrobot (*21*). As a result, materials that predictably degrade and disappear in a safe manner have become increasingly important for medical applications (*22*). Medical microrobots developed so far have not tackled the issue of biodegradability, so it complicates their clinical use due to possible adverse effects in the body.

Here, we report the design of 3D-printed, biodegradable microrobotic swimmer, which accomplishes its task of active therapeutic and diagnostic release based on the environmental sensing of matrix metalloproteinase 2 (MMP-2) enzyme. At the pathological concentrations of MMP-2, the microswimmer switches on an accelerated-drug release pathway by rapid swelling of its hydrogel network. Upon the complete breakdown of the microswimmer network, Anti-ErbB 2 antibody-tagged magnetic contrast agents are released into the local environment for targeted cell labeling, and thereby promising follow-up evaluation of the preceding therapeutic intervention. We describe the fabrication of a structurally complex, compositionally heterogeneous microswimmer using direct laser writing inside a nanocomposite of magnetic hydrogel precursor. The precursor comprises iron oxide nanoparticles dispersed in the gelatin methacryloyl, a photocrosslinkable semi-synthetic polymer derived from collagen. The bulk hydrogels of photocrosslinked gelatin methacryloyl were used in drug delivery and tissue engineering applications for its biocompatibility (*23*). Gelatin also contains target cleavage sites for MMP-2, thereby appealing as a biodegradable structural material for microrobots (*24*). The present study also shows hydrogels as a useful material platform for microswimmers to carry concentrated therapeutics in their bulk volume. Altogether, the findings of the present work represent a leap toward clinical medical mobile microrobots that are capable of sensing, responding to the local microenvironment, and performing specific diagnostic or therapeutic tasks using their smart composite material architectures in physiologically complex environments.

## RESULTS AND DISCUSSION

### Design, Fabrication and Mobility of Microswimmers

As the swimmer size goes to the microscopic scales, so the viscous forces begin to dominate over the inertial forces. As a result, a microswimmer needs to do continuous non-reciprocal motions to break spatial and temporal symmetries to generate a forward thrust (*25*). To comply with the same challenge, microorganisms in nature have evolved elaborate locomotion strategies, such as helical rotation of bacterial flagella, and the beating of paramecium cilia, which have so far inspired many synthetic swimmer designs (*26–31*). Inspired by a similar mechanism, the design of our microrobotic swimmer is illustrated in Fig. 1. From an empirical point of view, it comprises a cylindrical core wrapped by a double helix, and the cylinder has cones at both ends. Due to the chirality of the double helix, the rotational motion of the microswimmer is coupled to its translational motion. The basic geometry of the microswimmer primarily concerns with increasing the volume-to-surface ratio, with the goal of accommodating concentrated therapeutics in its bulk. Previous designs were limited to a simple helix, and the materials used to make them yielded non-porous architectures. As a result, such designs only concerned with the applications of cargo transport on the swimmer surface, which put significant limitations over the amount of the deliverable cargo, and hence the potential efficacy of the microrobotic operations (*29, 32, 33*).

**Fig. 1.**
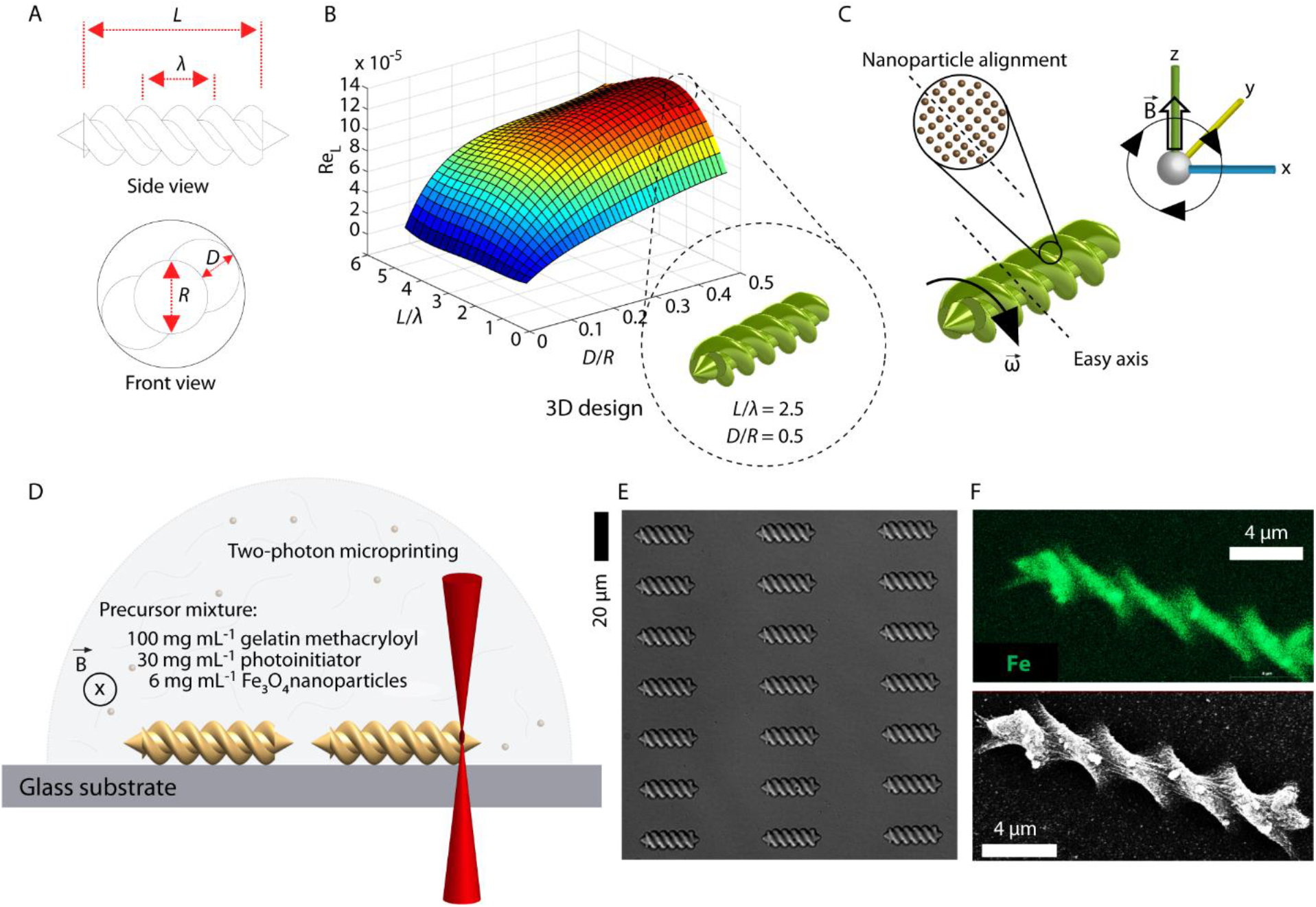
Design and 3D fabrication of biodegradable hydrogel microrobotic swimmers. (A) The empirical design of the double-helical microswimmer. (B) Computational fluid dynamics simulation for Reynolds number with respect to *L/λ* and *D/R* ratios, calculated for water at room temperature. The maximum forward swimming velocity was found with *L/λ* = 2.5 and *D/R* = 0.5 for the given design space sweep study. (C) Alignment of the magnetic nanoparticles that defines an easy axis normal to the helical axis, thereby allowing rotational motion under rotating magnetic fields. (D) 3D fabrication of the microswimmers using two-photon polymerization. During the fabrication process, a continuous magnetic field was applied to keep the nanoparticles aligned. (E) Optical microscope differential interference contrast (DIC) image of a microswimmer array. (F) Energy-dispersive spectroscopy (EDS) mapping of iron confirming the homogeneous embedding of the iron oxide magnetic nanoparticles inside the microswimmer body.

The dimensions of the microswimmer were determined by four parameters: *L*, λ, *D*, and *R* (Fig. 1a). The length, *L*, was fixed to 20.0 μm, and the diameter of the inner cylinder, *R*, to 3.3 μm, to have a microswimmer on the size order of an average mammalian cell. We then sought to maximize the swimming speed by varying the wavenumber, *L/λ*, and *D/R* in a series of computational fluid dynamics (CFD) simulations and built the equation of motion for the microswimmer (Supplementary Fig. S1-S3. Also see Supplementary Text S1 for a detailed discussion). Rotation-translation coupling was observed to lie in an optimum region, yielding the maximum swimming velocity within the confines of the studied design space (Fig. 1b and Supplementary Fig. S4). The coupling decreased toward the long and short wavelengths, as, in both cases, the structure geometry lost its chirality and converged to a cylinder, and hence lost its ability to generate forward thrust. The coupling also decreased by decreasing the cord radius, *D*, of the double helix, because the interaction surface of the double helix with the surrounding fluid went smaller. Altogether, the dimensionless ratios of *L/λ*=2.5 and *D/R*=0.5 were evaluated to give the highest swimming speed given the design considerations. Concerning the trajectory stability along a single line, a single helix is expected to swim with lower stability at non-integer wavenumbers, here 2.5, due to the balancing of the hydrodynamic forces throughout the helices (*34*). Nevertheless, the design in the form of double helix compensates the trajectory stability. The lateral drift was found to be an order of magnitude higher with the single-helix configuration than that of the double helix configuration (Supplementary Fig. S5). The wobbling velocity, *i.e*., the lateral angular velocities of the microswimmer with single helix configuration, was also an order of magnitude higher with the single-helix (Supplementary Fig. S6). These simulations showed that the double helix represents an advantageous design scheme in favor of hydrodynamic efficiency, and thus power requirements.

The use of magnetic fields is a prominent way of remote powering and control of medical microrobots (*35*). In contrast to other untethered power transfer alternatives, such as light and chemical signals, magnetic fields are able to safely penetrate biological tissues and other materials (*12*). To rotate the double helix, rotating magnetic fields are needed, which then create a torque on the microswimmer through a magnetic axis defined perpendicular to the helical axis. Assuming the external field is invariant, the magnetic force scales with the volume of the magnetic material, *i.e*., *L*^3^, while the equivalent force from the magnetic torque scaling with *L*^2^. Thus, swimming by magnetic torque-induced rotation is preferable at smaller scales, *ca*. less than 100 μm, due to its higher efficiency of thrust generation (*20*). We sought to embed superparamagnetic iron oxide nanoparticles in the form of a nanocomposite to impart magnetizability to the microswimmer body. Embedding iron oxide nanoparticles has certain advantages over magnetic surface coating, such as with cobalt or nickel. Cobalt and nickel are considerably toxic to the living environment, whereas iron oxide nanoparticles are mostly regarded biofriendly (*36*). Coating the robot surface would also drastically reduce the availability of the microswimmer bulk for cargo loading in and release from the porous hydrogel network. Biofunctionalized magnetic nanoparticles could also be exploited as smart contrast agents for targeted cell labeling toward medical imaging and therapy (*37*). In order to create an effective magnetic torque for rotation, introduction of a magnetization axis perpendicular to the helical axis was necessary, which was accomplished by the alignment and entrapment of the magnetic nanoparticles under uniform magnetic field during the fabrication process (Fig. 1c, d) (*38*).

To realize such a design with highly demanding geometrical and compositional aspects, we employed 3D-microprinting based on two-photon polymerization, or direct laser writing. Briefly, this fabrication technique relies on the simultaneous absorption of two half-energy photons by the photoinitiator at the tight focal spot of a femtosecond-pulsed infrared laser light. As the two-photon absorption occurs only in a fraction of the focal volume, and the polymer precursor is transparent to the infrared laser wavelength, *i.e*., 780 nm, spatiotemporal control of the two-photon absorption enables highly complex 3D computer-aided designs with sub-micron features. For the printing, we prepared a nanocomposite hydrogel precursor containing 100 mg mL^-1^ gelatin methacryloyl, 30 mg mL^-1^ photoinitiator, and 6 mg mL^-1^ iron oxide nanoparticles, which we ensured to be compatible with two-photon polymerization (Fig. 1d and Supplementary Fig. S7 and S8). The structural fidelity of the resulting microswimmers was in agreement with the design, and we did not observe undesired shrinkage or swelling in the polymerization and development steps (Fig. 1e). Energy-dispersive x-ray analysis proved homogenous distribution of the iron oxide nanoparticles inside the microswimmer (Fig. 1f and Supplementary Fig. S9).

The total amount of magnetic nanoparticles loaded in the mesh network of the microswimmer body is critical for the magnitude of magnetization, and hence the swimming speed. Decreasing the amount will result in lower step-out frequencies, thereby reducing the achievable swimming speed. Therefore, we aimed to maximize the volume fraction of the magnetic nanoparticles in the precursor suspension. Nevertheless, increased concentrations of nanoparticles from 0.1 to 10 mg mL^-1^ narrowed down the effective laser energy density range for both low and high laser intensities (Fig. S10). At lower laser intensities, the aggregates of nanoparticles at higher nanoparticle concentrations physically blocked the propagation of the polymerization because there are fewer chains generated to complete the assigned laser trajectories, and insufficiently linked structures lowered the structural quality. At higher laser intensities, nanoparticle aggregates strongly interacted with the laser light, resulting in local heating, bubble generation, and structural damage. This problem could be potentially circumvented with iron oxide nanoparticles that have better colloidal stability, and hence they have lesser tendency for aggregation at higher concentrations. In the current nanoparticle design, microscopic aggregates were evident during polymerization above 6 mg mL^-1^ iron oxide nanoparticle concentration. In this regard, this threshold concentration was used in the rest of the microswimmer preparations, as they were homogeneously dispersed in the solution without any particle aggregation or agglomeration (Supplementary Fig. S8). Additionally, the two-photon lithography method does not reach temperatures that would affect the magnetic properties of the iron oxide nanoparticles.

For magnetic mobilization, we utilized a custom six-coil electromagnetic setup to create rotating magnetic fields that exert a computer-controlled magnetic torque on the microswimmer’s long axis (Supplementary Fig. S3 and S11). We determined the step-out frequency of the microswimmers loaded with 6 mg mL^-1^ iron oxide nanoparticles by increasing input frequency at every 10 seconds starting from 1 Hz to 6 Hz with 1 Hz step size at 20 mT. The swimming velocity increased linearly with increasing excitation frequency (Fig. 2a and Supplementary Movie S1). The microswimmers exhibited predominantly wobbling behavior in the frequency range of 1-3 Hz. The cork-screw motion was observed in the range of 3-5 Hz. The step-out frequency of the microswimmers was 6 Hz. We characterized the long-distance trajectory of the microswimmers by actuating them at 5 Hz, just below their step-out frequency, to determine their average swimming speed (3.36 ± 0.71 μm s^-1^) (Fig. 2b and Supplementary Movie S2). Temporally constant rotating fields in the work space allow navigation of microswimmers as small teams (Supplementary Movie 2).

**Fig. 2:**
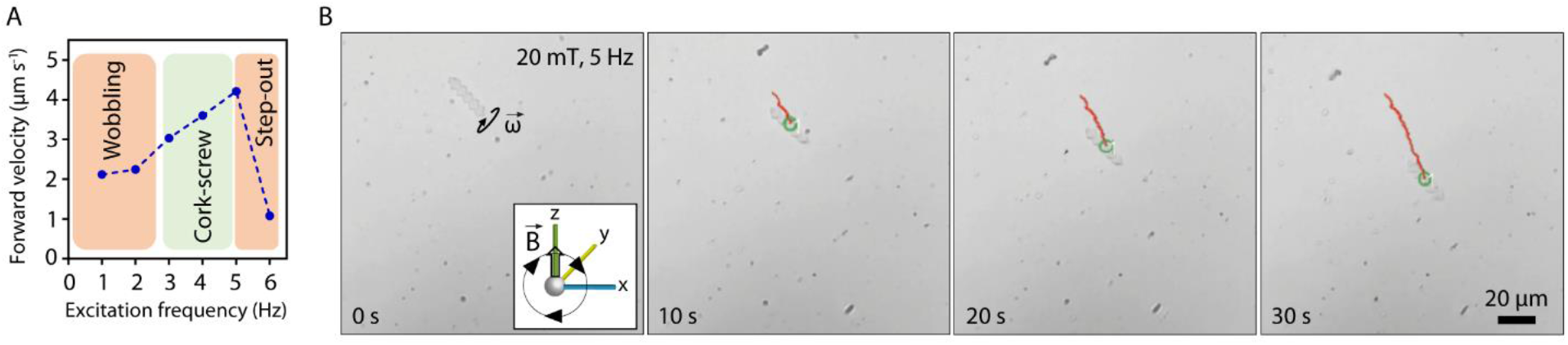
Swimming trajectory of a double-helix microswimmer under a rotating magnetic field. (A) The step-out frequency of the microswimmers containing 6 mg mL^-1^ iron oxide nanoparticles was found to be around 6 Hz. (B) Image sequence of the hydrogel microswimmer actuated under a rotating magnetic field with the magnetic field strength and excitation frequency being 20 mT and 5 Hz, respectively. The average velocity of the microswimmers was found to be 3.36 ± 0.71 μm s^-1^.

Above swimming motility simulations and experiments were conducted in water at room temperature for an easier quantitative first study. However, since future *in vivo* experiments need to be conducted in biological fluids, which are non-Newtonian and heterogeneous, we will simulate and test the proposed microswimmer in body fluids at body temperature in the future.

### Biodegradability with MMP-2

We explored the biodegradability of the microswimmers with MMP-2 enzyme, also known as gelatinase A (*39*). MMPs operate in the extracellular environment and play an important role in tissue remodeling by degrading various extracellular matrix components. These enzymes could, therefore, represent an appealing target for the proteolytic degradation of our microswimmers once they have completed their prescribed tasks. In a healthy individual, this enzyme is reportedly present at various concentrations, yet typically in the range of 140-200 ng mL^-1^, depending on type of the tissue it is expressed (*40, 41*). At an unrealistically high concentration of MMP-2, *i.e*., 100 μg mL^-1^, we observed that the microswimmers were entirely degraded within an hour *in vitro* at 37 °C (Supplementary Movie 3). The complete degradation time was a function of the initial MMP-2 concentration (Figs. 3a,b). At more realistic concentrations, such as 4 μg mL^-1^, it took around 5 h for complete degradation, 67 h at 0.500 μg mL^-1^, and 118 h at 0.125 μg mL^-1^, *i.e*., the healthy physiological level. Very recently, salt-based erosion of inorganic microswimmers was shown for the route of microswimmer degradation in the living environment (*42*). To the best of our knowledge, though, the present study is the first example of a microrobotic swimmer that is completely removed by an enzymatic process, which leaves no detectable toxic residues behind (Figs. 3d, e). While we did not observe an acute toxicity with the SKBR3 cell line, even in the presence of high iron oxide nanoparticle concentrations, given the potential long-term toxicity risks raised by the use of nanoparticles should be taken into account in the line of the current literature (*36*). We anticipate that the degradation time could be further extended by introducing non-degradable methacryloyl polymers, such as poly(ethylene glycol) in the microswimmer network, so that the density of the enzyme-recognition sites would be lowered.

**Fig. 3.**
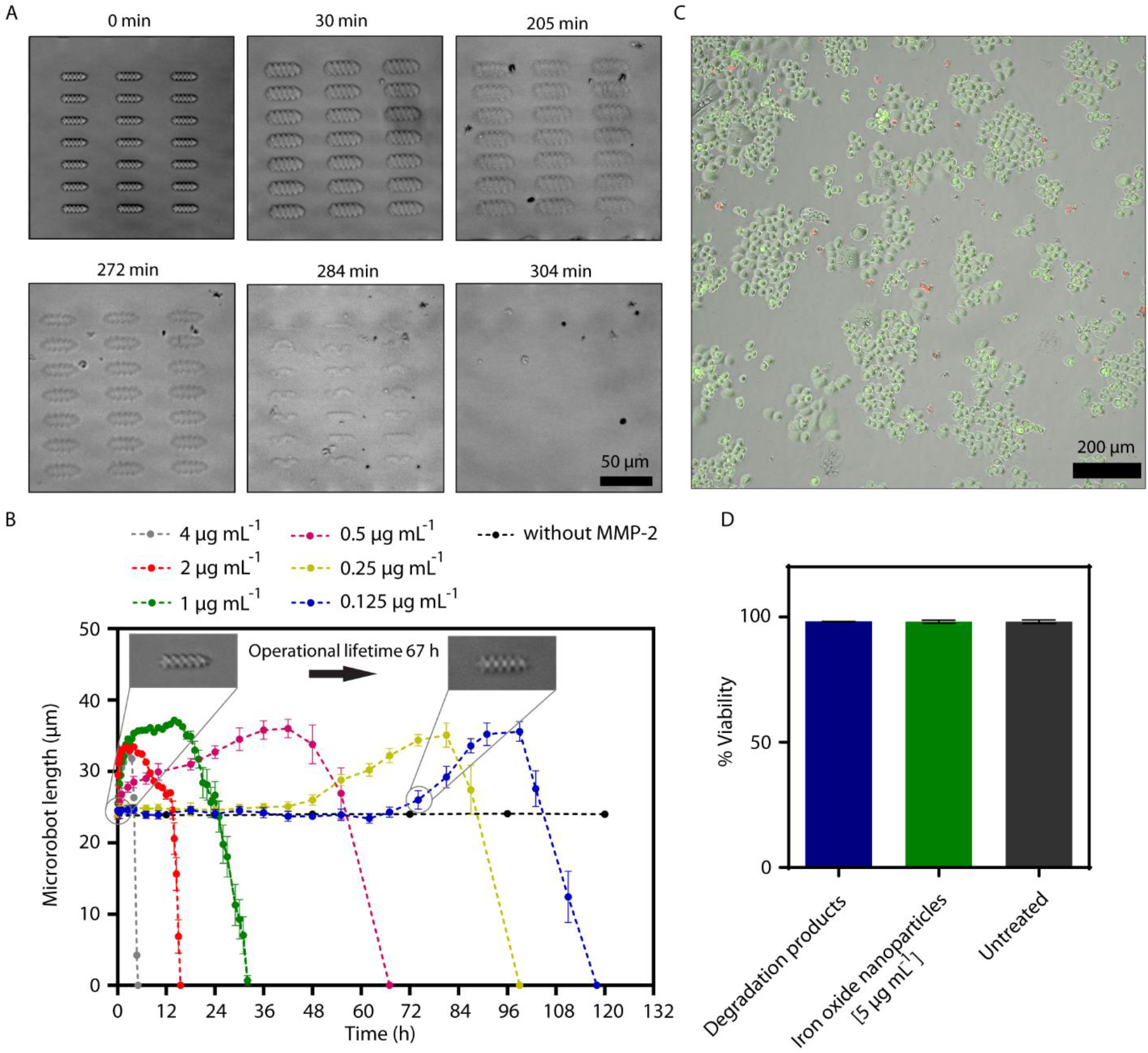
Biodegradation of the hydrogel microswimmers by the MMP-2 enzyme. (A) DIC images of a degrading microswimmer array in the presence of 4 μg mL^-1^ enzyme. Degradation starts with the rapid swell of the microswimmers followed by the collapse of the entire network. (B) Enzymatic degradation of the microswimmers. At the physiological level, MMP-2 degrades the microswimmers within 118 h. Enzymatic susceptibility introduces a new concept of operational lifetime, defined as the time period that microswimmer preserves its original morphology for proper navigation. The operational lifetime is a function of the enzyme concentration in the microenvironment. Data are presented as mean ± standard deviation. (C) Live (green) and dead (red) SKBR3 breast cancer cells treated with the degradation products of the microswimmers. (D) Quantitative analysis of the acute toxicity induced by the degradation products of the microswimmers in comparison with 5 μg mL^-1^ iron oxide nanoparticles and untreated cells. Data are presented as mean ± standard deviation. (P > 0.05, one-way ANOVA)

A ubiquitous response pattern of the microswimmer to the enzyme emerged in the degradation process. Shortly after the introduction of MMP-2 above the physiologically relevant concentration, we observed rapid swelling of the microswimmer body up to three folds of its initial volume, followed by the collapse of the entire hydrogel network over time (Fig. 3b). Considering the relationship between the rate of water diffusion into the polymer network and the kinetics of the bond cleavage, the observed phenomenon is reminiscent of homogenous, or bulk, erosion. In bulk degradation, the loss of mass also occurs throughout the material in a homogeneous manner (*22*). As the rate of water diffusion exceeds the rate of the bond cleavage reaction, uniform swelling is observed long before the collapse of the network (*43*). Taking into account of the morphological change of the microswimmers, a therapeutic time window appears for the microrobotic tasks to be carried out in a controlled way. The operational lifetime ends before the complete degradation, when the microswimmers start swelling (a volumetric expansion above 2% was considered as the threshold value), because the microswimmers would then lose their ability to navigate due to significant loss of the structural chirality. The operational lifetime of the microswimmers in the physiological environment is around 67 h while the complete degradation time is 118 h. As a result, the task of a biodegradable microrobot concerning its navigation and localization should be accomplished within the operational lifetime (Fig. 3b).

### Boosted Drug Release

A major compositional fraction of hydrogels is water, typically accounting for more than 90% by mass. Highly porous hydrogel networks can thus very effectively sequester high amounts of therapeutic and diagnostic cargo types, which can then be released in response to various input signals with controllable spatial and temporal schemes (*44, 45*). Overexpression of matrix metalloproteinases is associated with almost any cancer types, because degradation of the matrix promotes tumor-cell growth, migration, invasion, metastasis, and angiogenesis (*39, 46*). For example, the concentration of MMP-2 above 200 ng mL^-1^ in the serum was associated with colorectal cancer patients whereas the quantitative amounts are largely unknown for other cancer types. MMP-2 could thus represent a valuable biomarker for the disease state of the tissue to be sensed and acted upon by the microswimmers. In this regard, the rapid swelling of the microswimmers as an initial response at the elevated concentrations of MMP-2 could serve as a switch for accelerated drug release. As a hydrogel swells, the mesh size increases, and the entrapped drug is released (*45*). Since the mesh size is correlated with the extent of swelling, the initial MMP-2 concentration could regulate the overall release kinetics. To this end, we first studied the relationship between the swelling kinetics and the initial enzyme concentration (Fig. 4a). At 0.125 μg mL^-1^ MMP-2 concentration, around the physiological level, swelling response was not evident in the first 30 min. At 0.250 μg mL^-1^, the swelling began to occur after 20 min following the introduction of the enzyme. At 0.5 and 1 μg mL^-1^, the swelling started as soon as the enzyme was introduced. A typical limitation of swelling-controlled drug delivering hydrogels is their slow responsiveness due to the slow diffusion of water. However, as our microswimmer is small, so is the diffusion length, and is highly porous, the response was fast and robust. As a result, the volume of the microswimmers increased by *ca*. 60 *%* within the first 30 min in the presence of 1 μg mL^-1^ MMP-2. This volumetric expansion considerably accelerated, (more than 20%) the model drug-analog dye release as an initial response to high enzyme concentration (Fig. 4b and Fig. S12). Control over the drug release kinetics by means of the cues in the pathological microenvironment is particularly valuable for effective and proportional engagement with the disease. In this regard, the present work is the first to address encoding functional properties to microswimmers for responding the changes in the microenvironment. Biodegradation is highly valuable in order to be able to fully utilize the drug cargo in the hydrogel network. Without degradation, almost the half of the payload remains inside the network, and hence is a sign of poor bioavailability. On the other hand, network degradation increases the bioavailability of the drug payload by releasing all of its content from the microswimmers (Figs. 4b,c).

**Fig. 4.**
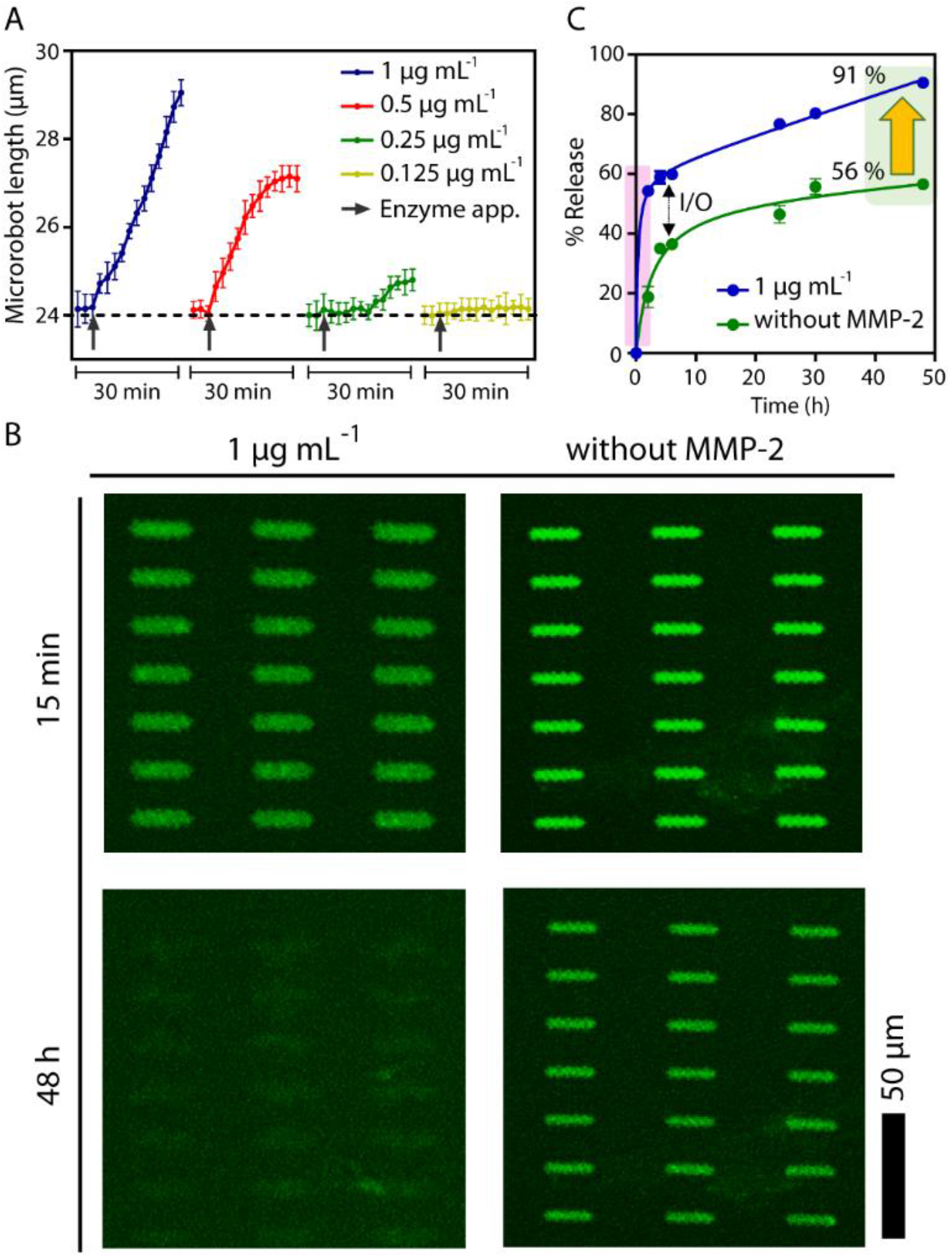
Enzymatically controlled drug release from the hydrogel microswimmers. (A) Elevated concentrations of MMP-2 cause rapid swelling of the microswimmer body, thereby acting as a switch for accelerated drug release. Data are presented as mean ± standard deviation. (B) At 1 μg mL^-1^ MMP-2 concentration, accelerated drug release in the first few hours (the pale pink region) is attributed to swelling-mediated mesh size increase. At the end of two days, almost all the payload is released from the degraded microswimmers whereas half of the content is retained in the non-degraded one, which severely reduces the bioavailability of a significant portion of the drug to be delivered (pale green region). Data as presented with mean ± standard deviation. (C) Epifluorescence images of microswimmers with loaded dextran-FITC cargo used as a model macromolecular drug equivalent. Enhanced drug bioavailability is evidenced by the enzymatic degradation of the network, which releases its entire content.

### Targeted Cell Labeling

The enzymatic collapse of the hydrogel network further releases the magnetic nanoparticles that are used to provide the magnetic torque for locomotion. We envision that targeted labeling of tumor cells with these nanoparticles could enable follow-up evaluation of the intervened tissue sites in a future clinical scenario. To this end, we modified the nanoparticle surface to display anti-ErbB 2 antibody for targeting ERBB2 receptors, which is overexpressed in the breast cancer cell line SKBR3 (Fig. 5a). For the ease of analysis *in vitro*, the nanoparticles were also modified with fluorophores. The antibody-tagged nanoparticles were embedded inside the microswimmers similarly to their non-modified counterparts (Fig. 5b). Following the degradation of the microswimmers, the functional nanoparticles released into the environment could label 39.3% of SKBR3 cells (Fig. 5c and Fig. S13). On the other hand, only 9.5% of the cell population was labelled without the surface modification of the nanoparticles, which shows a small degree of nonspecific interaction existed between the cell and nanoparticles, as well (Fig. 5d). This is the very first proof of the concept to encode a combination of therapeutic and diagnostic control scenarios in a microrobotic operation, where the treatment by means of drug release is monitored by mapping the remaining target cells with the magnetic contrast agents. Complete biodegradation enables the release of internally embedded medical contrast agents. Antibody-modified contrast agents diffuse around to target the untreated tissue sites, which could be monitored to assess the therapeutic efficiency of the treatment, and to identify new intervention sites using imaging techniques, such as magnetic resonance imaging (Fig. S14).

**Fig. 5.**
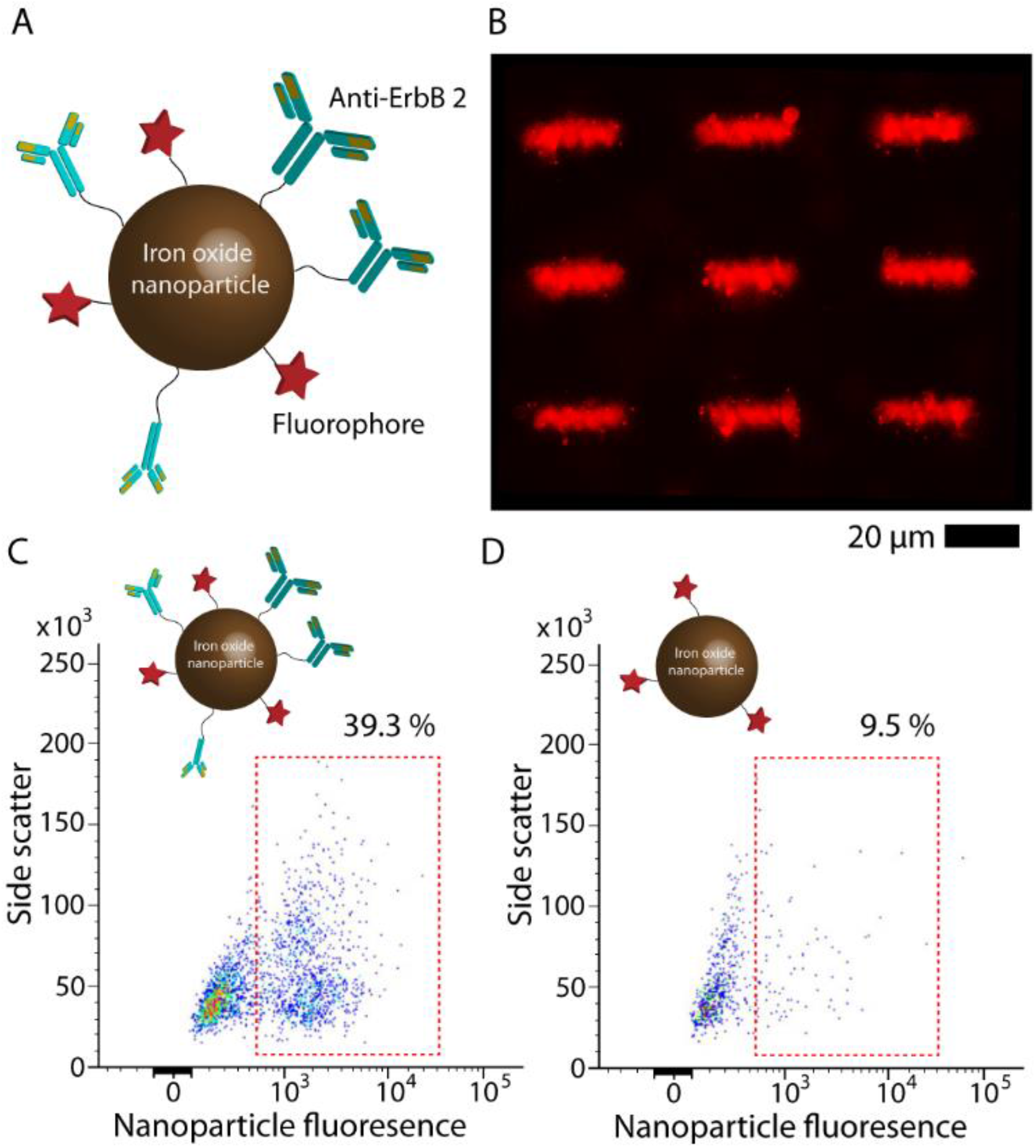
Targeted cell labeling with the magnetic nanoparticles released from collapsed microswimmers towards diagnostic *in vivo* imaging. (A) The design of superparamagnetic iron oxide nanoparticles of 50 nm size functionalized with a fluorophore and anti-ErbB 2 antibody for targeted labeling of ERBB2-overexpressing breast cancer SKBR3 cells. (B) Epifluorescence image of microswimmers embedded with the nanoparticles. (C) Targeted labeling of SKBR3 with the anti-ErbB 2 modified magnetic nanoparticles released upon the MMP-2-mediated degradation of the microswimmers. In the absence of anti-ErbB 2, the nanoparticles fail to target SKBR3 cells (D).

## CONCLUSION

As the clinical interest of robotic devices is shifting to the development of small, autonomous, or remotely controlled systems, challenges remain regarding material biocompatibility, biodegradability, and execution of functional tasks in a programmed way. An ideal material solution should convey the idea of short-term inertness in the body while a microrobot should be degradable with the lowest possible waste profile in the long-term. Here, we designed, explored, and characterized *in vitro* a hydrogel based biodegradable helical microswimmer remotely controlled by rotating magnetic fields. Owing to their emergent physical properties and capability to protect labile drugs from degradation, hydrogels could be programmed for various physiochemical interactions with the encapsulated drugs to control drug release. The use of a biopolymer derivative, gelatin methacryloyl, to make microrobots will pave the way for making patient-specific microrobots using their own biomaterials. Such a personalized strategy could largely circumvent potential concerns of the immunogenicity. Over the past decade, a variety of microswimmers has been proposed; however, here we complementarily concentrated on the sensing and responding the changes in the medical environment, such as a disease marker enzyme, MMP-2, which triggered the microswimmer to accelerate the drug release at the tumor site. Active and targeted delivery of multifunctional cargo types, such as drugs, imaging agents, genes, and RNA, are the major objectives of microrobotic operations in order to make a realistic impact in the near future. While such capabilities could enable high impact applications in targeted delivery, microsurgery, tissue engineering, and regenerative medicine; in the longer term, they could aspire new treatment models for genetic diseases by single cell-level proteins or nucleic acid delivery and roaming the body for disease prediction and prevention.

## MATERIALS AND METHODS

### Preparation of 3D-printable superparamagnetic magnetic precursor

100 mg mL^-1^ gelatin methacryloyl, 30 mg mL^-1^ lithium phenyl(2,4,6-trimethylbenzoyl) phosphinate, 6 mg mL^-1^ iron oxide nanoparticles, coated with poly(ethylene glycol) amine, of 50 nm hydrodynamic size (Chemicell GmbH, Germany), were mixed in ultrapure water with vortex mixing and ultrasound sonication. The resulting suspension was dropped on a glass slide, on which the fabrication was carried out. Commercially available Direct Laser Writing system (Photonic Professional, Nanoscribe GmbH, Germany) with a 63x oil-immersion objective (NA 1.4) was used for 3D printing of the microrobots via two-photon polymerization. Laser power and galvanometric mirror x- and y-scanning speeds were optimized for printing as 23.5 mW and 3.0 x 10^5^μm s^-1^, respectively (Supplementary Fig. S10). The overall microprinting rate was measured to be 10 s for a single microrobot. DIC images were taken with Nikon Eclipse Ti-E inverted microscope. EDS measurements were made with a Zeiss Gemini 500 scanning electron microscope equipped with Quantax EDS (Bruker).

Microswimmers printed on glass substrate, and kept at 4°C refrigerator overnight in water, were equilibrated to room temperature prior to recombinant human MMP-2 (Sigma Aldrich) addition at various concentrations (Fig. 3b and Fig 4a.). Time lapse images were taken using a Nikon Eclipse Ti-E inverted microscope at 20x magnification in the DIC mode. The enzyme solutions incubated with the microswimmers were replenished every 12 h in order to minimize the impact of enzymatic deactivation at long time periods. For analysis, the length of the microswimmers (21 microswimmers for each time frame) was measured using Nikon NIS AR Element analysis software.

In order to investigate the toxicity of the microswimmer degradation products, the viability of ERBB2-overexpressing SKBR3 cells (DSMZ, Germany) was tested by Live/Dead cell imaging kit (Invitrogen) after 24 h of treatment with degradation products (from 2349 microswimmers degraded in 2 μg mL^-1^ MMP-2) and bare nanoparticles, according to manufacturer’s instructions. SKBR3 cells were cultured in McCoy’s 5a Modified Medium supplemented with 10% FBS, penicillin (50 UI ml^-1^) and streptomycin (50 μg ml^-1^). Cells were grown at 37 °C and 5% CO2 in a humidified environment and sub-cultured before confluence using trypsin/EDTA. Cells with no treatment was taken as a control. Live and dead cells were observed under fluorescent microscope using FITC and TRITC filters.

### Drug release assay

Microswimmers printed on glass substrate were loaded with a drug-equivalent macromolecule, dextran-FITC (Mw Sigma Aldrich), by immersing the glass into 1 mM dextran-FITC (Mw 10,000 Da)-containing aqueous solution overnight at 4°C. The microswimmers were equilibrated to room temperate, and washed with copious amount of water for 10 min before the drug release measurements. Drug release from microswimmers was analyzed using a spinning disc (Yokogawa, Japan) confocal microscope (Nikon Eclipse Ti-E). Before the experiment, a separate array of microswimmer was exposed to the varying intensities of laser light and exposure times in order to eliminate the bleaching effect (Fig. S12). Confocal fluorescence images were then acquired from the actual release samples every hour for a period of 12 h, then every 6 h by the end of 48 h. MMP-2 was added only in the beginning, and no enzyme replenishment was done afterwards. Fluorescence intensities over 21 microswimmers were measured using Nikon software. Background fluorescence was subtracted from the measured values.

### Cellular viability and targeted cell labeling

For cell labeling assay, SKBR3 cells at 1×10^5^ cells/ml density were incubated with microswimmer degradation products (from 2349 microswimmers degraded in 2 μg mL^-1^ MMP-2) in HEPES buffered saline for 1 h at 37 °C. Nanoparticles without anti-ERBB2 antibody served as the negative control. After incubation, cells were centrifuged, washed and resuspended in buffer and analyzed by flow cytometer (BD FACSMelody™). Untreated SKBR3 cells were used to set the gates and a total of 5000 events were acquired for each analysis after gating on singlets (Fig. S13).

### Statistical analysis

All experiments were independently repeated at least four times. In each repeat, at least 21 individual microswimmers were counted. The error bars represent ± standard deviation. The statistical analyses were done with one-way analysis of variance (ANOVA). A *P* value higher than 0.05 was considered statistically significant. The quantitative results represent the overall average of independent and technical replicas.

## SUPPLEMENTARY MATERIALS

Text S1. Hydrodynamic optimization of the microswimmer body.

Text S2. Materials and methods.

Text S3. Building the magnetic coil setup.

Fig. S1. The numerical setup with the boundary conditions.

Fig. S2. The response surface of c6z with respect to the ratio of cord-radius to core diameter, *D/R*, vs. the ratio of core length to the wave length, *L/λ*.

Fig. S3. The design of the air-core Helmholtz coils.

Fig. S4. The ratio of the displacement by forward-swimming velocity, *Uz*, per unit time to the body length of the robot, *L*.

Fig. S5. The lateral drift of the microswimmers with different helical arrangements.

Fig. S6. Wobbling motion of the robot, *i.e*., lateral rigid-body rotations, with respect to different helix arrangement.

Fig. S7. Synthesis and characterization of gelatin methacryloyl.

Fig. S8. Preparation of superparamagnetic hydrogel precursor solution.

Fig. S9. Confirmation of iron inside microswimmers with energy-dispersive x-ray spectroscopy. Fig. S10. Optimization of the femtosecond laser energy density for the maximized structural fidelity and iron oxide nanoparticle density inside microrobot.

Fig. S11. The assembled magnetic coil pairs placed under the stereomicroscope.

Fig. S12. Confocal fluorescence imaging optimizations and Dextran-FITC loaded microswimmers at t=0 min.

Fig. S13. Targeted labeling of SKBR3 breast cancer cells with anti-ErbB 2 antibody-modified magnetic nanoparticles.

Fig. S14. Application scenario of the 3D-printed, biodegradable microrobotic swimmers in a clinical setting.

Movie S1. Determination of the step-out frequency of the biodegradable microswimmers.

Movie S2. Long-distance trajectory of the biodegradable microswimmers.

Movie S3. Enzymatic degradation of the microswimmers by MMP-2.

References (47-60)

## Acknowledgements

The authors acknowledge funding from the Max Planck Society and the Max Planck ETH Center for Learning Systems.

## REFERENCES AND NOTES

1. A. Darzi, Y. Munz, The impact of minimally invasive surgical techniques. Annu. Rev. Med., 55, 223–237 (2004).

2. B. Davies, Robotic surgery – A personal view of the past, present and future. Int. J. Adv. Robot. Syst., 12, 54 (2015).

3. M. Sitti, Miniature devices: Voyage of the microrobots. Nature, 458, 1121–1122 (2009).

4. B. J. Nelson, I. K. Kaliakatsos, J. J. Abbott, Microrobots for minimally invasive medicine. Annu. Rev. Biomed. Eng., 12, 55–85 (2010).

5. M. Sitti, H. Ceylan, W. Hu, J. Giltinan, M. Turan, S. Yim, E. Diller, Biomedical applications of untethered mobile milli/microrobots. Proc. IEEE, 103, 205–224 (2015).

6. S. Fusco, F. Ullrich, J. Pokki, G. Chatzipirpiridis, B. Özkale, K. M. Sivaraman, O. Ergeneman, S. Pané, B. Nelson, Microrobots: a new era in ocular drug delivery. Expert Opin. Drug Deliv., 11, 1815–1826 (2014).

7. S. Martel, Beyond Imaging: Macro-and microscale medical robots actuated by clinical MRI scanners. Sci. Robot., 2, eaam8119 (2017).

8. X. Wang, M. Luo, H. Wu, Z. Zhang, J. Liu, Z. Xu, W. Johnson, Y. Sun, A three-dimensional magnetic tweezer system for intraembryonic navigation and measurement. IEEE Trans. Robot., 34, 240–247 (2018).

9. M. P. Kummer, J. J. Abbott, B. E. Kratochvil, R. Borer, A. Sengul, B. J. Nelson, OctoMag: An electromagnetic system for 5-dof wireless micromanipulation. IEEE Trans. Robot., 26, 1006–1017 (2010).

10. S. Jancik, D. Houle, M. Lafleur, L. Gaboury, M. Tabrizian, N. Kaou, M. Atkin, T. Vuong, G. Batist, N. Beauchemin, D. Radzioch, S. Martel, Magneto-aerotactic bacteria deliver drug-containing nanoliposomes to tumour hypoxic regions. Nat. Nano., 11, 941–947 (2016).

11. B. E.-F. de Ávila, P. Angsantikul, J. Li, M. A. Lopez-Ramirez, D. E. Ramírez-Herrera, S. Thamphiwatana, C. Chen, J. Delezuk, R. Samakapiruk, V. Ramez, M. Obonyo, L. Zhang, J. Wang, Micromotor-enabled active drug delivery for in vivo treatment of stomach infection. Nat. Commun., 8, 272 (2017).

12. H. Ceylan, J. Giltinan, K. Kozielski, M. Sitti, Mobile microrobots for bioengineering applications. Lab Chip, 17, 1705–1724 (2017).

13. J. Li, B. Esteban-Fernández de Ávila, W. Gao, L. Zhang, J. Wang, Micro/nanorobots for biomedicine: Delivery, surgery, sensing, and detoxification. Sci. Robot, 2, eaam6431 (2017).

14. D. Chu, X. Dong, Q. Zhao, J. Gu, Z. Wang, Photosensitization priming of tumor microenvironments improves delivery of nanotherapeutics via neutrophil infiltration. Adv. Mater., 29, 1701021 (2017).

15. J. Cohen, IL-12 Deaths: Explanation and a Puzzle. Science, 270, 908–908 (1995).

16. G. K. Such, Y. Yan, A. P. R. Johnston, S. T. Gunawan, F. Caruso, Interfacing materials science and biology for drug carrier design. Adv. Mater., 27, 2278–2297 (2015).

17. B. Siciliano, O. Khatib, Springer Handbook of Robotics. (Springer-Verlag New York, Inc., 2007).

18. C. R. Reid, H. MacDonald, R. P. Mann, J. A. R. Marshall, T. Latty, S. Garnier, Decision-making without a brain: how an amoeboid organism solves the two-armed bandit. J. R. Soc., Interface, 13, 20160030 (2016).

19. H. Ceylan, I. C. Yasa, M. Sitti, 3D chemical patterning of micromaterials for encoded functionality. Adv. Mat., 29, 1605072 (2017).

20. M. Sitti, Mobile Microrobotics. (MIT Press, ed. 1, 2017).

21. B. D. Ulery, L. S. Nair, C. T. Laurencin, Biomedical applications of biodegradable polymers. J. Polym. Sci., Part B: Polym. Phys., 49, 832–864 (2011).

22. S. Lyu, R. Sparer, D. Untereker, Analytical solutions to mathematical models of the surface and bulk erosion of solid polymers. J. Polym. Sci., Part B: Polym. Phys., 43, 383–397 (2005).

23. K. Yue, G. Trujillo-de Santiago, M.M. Alvarez, A. Tamayol, N. Annabi, A. Khademhosseini, Synthesis, properties, and biomedical applications of gelatin methacryloyl (GelMA) hydrogels. Biomaterials, 73, 254–271 (2015).

24. M. Toth, A. Sohail, R. Fridman, in Metastasis Research Protocols, M. Dwek, S. A. Brooks, U. Schumacher, Eds. (Humana Press, Totowa, NJ, 2012), pp. 121–135.

25. E. M. Purcell, Life at low Reynolds number. Am. J. Phys., 45, 3–11 (1977).

26. T. Honda, K. I. Arai, K. Ishiyama, Micro swimming mechanisms propelled by external magnetic fields. IEEE Trans. Magn., 32, 5085–5087 (1996).

27. D. J. Bell, S. Leutenegger, K. M. Hammar, L. X. Dong, B. J. Nelson, in Proceedings IEEE International Conference on Robotics and Automation, 10 to 14 April 2007, Roma, Italy, pp. 1128–1133.

28. A. Ghosh, P. Fischer, Controlled propulsion of artificial magnetic nanostructured propellers. Nano Lett., 9, 2243–2245 (2009).

29. S. Tottori, L. Zhang, F. Qiu, K.K. Krawczyk, A. Franco-Obregón, B.J. Nelson, Magnetic helical micromachines: Fabrication, controlled swimming, and cargo transport. Adv. Mater., 24, 811–816 (2012).

30. P.L. Venugopalan, R. Sai, Y. Chandorkar, B. Basu, S. Shivashankar, A. Ghosh, Conformal Cytocompatible Ferrite Coatings Facilitate the Realization of a Nanovoyager in Human Blood. Nano Lett., 14, 1968–1975 (2014).

31. G.Z. Lum, Z. Ye, X. Dong, H. Marvi, O. Erin, W. Hu, M. Sitti, Shape-programmable magnetic soft matter. Proc. Natl. Acad. Sci. U. S. A., 113, E6007–E6015 (2016).

32. F. Qiu, S. Fujita, R. Mhanna, L. Zhang, B.R. Simona, B.J. Nelson, Magnetic helical microswimmers functionalized with lipoplexes for targeted gene delivery. Adv. Funct. Mater., 25, 1666–1671 (2015).

33. D. Walker, B. T. Käsdorf, H.-H. Jeong, O. Lieleg, P. Fischer, Enzymatically active biomimetic micropropellers for the penetration of mucin gels. Sci. Adv., 1, e1500501 (2015).

34. A. F. Tabak, S. Yesilyurt, Computationally-validated surrogate models for optimal geometric design of bio-inspired swimming robots: Helical swimmers. Comput. Fluids, 99, 190–198 (2014).

35. F. Qiu, B. J. Nelson, Magnetic Helical Micro-and Nanorobots: Toward their biomedical applications. Engineering, 1, 021–026 (2015).

36. M. Mahmoudi, H. Hofmann, B. Rothen-Rutishauser, A. Petri-Fink, Assessing the in vitro and in vivo toxicity of superparamagnetic iron oxide nanoparticles. Chem. Rev., 112, 2323–2338 (2012).

37. X.-H. Peng, X. Qian, H. Mao, A.Y. Wang, Z. Chen, S. Nie, D.M. Shin, Targeted magnetic iron oxide nanoparticles for tumor imaging and therapy. Int. J. Nanomed., 3, 311–321 (2008).

38. J. Kim, S.E. Chung, S.-E. Choi, H. Lee, H. J. Kim, S. Kwon, Programming magnetic anisotropy in polymeric microactuators. Nat. Mater., 10, 747–752 (2011).

39. M. Egeblad, Z. Werb, New functions for the matrix metalloproteinases in cancer progression., 2, 161 (2002).

40. K. Thrailkill, G. Cockrell, P. Simpson, C. Moreau, J. Fowlkes, R.C. Bunn, Physiological matrix metalloproteinase (MMP) concentrations: comparison of serum and plasma specimen. Clin. Chem. Lab. Med., 44, 503–504 (2006).

41. M. Groblewska, B. Mroczko, M. Gryko, A. Pryczynicz, K. Guzińska-Ustymowicz, B. Kędra, A. Kemona, M. Szmitkowski, M., Serum levels and tissue expression of matrix metalloproteinase 2 (MMP-2) and tissue inhibitor of metalloproteinases 2 (TIMP-2) in colorectal cancer patients. Tumor Biol., 35, 3793–3802 (2014).

42. X. Yan, Q. Zhou, M. Vincent, Y. Deng, J. Yu, J. Xu, T. Xu, T. Tang, L. Bian, Y.-X. J. Wang, K. Kostarelos, L. Zhang, Multifunctional biohybrid magnetite microrobots for imaging-guided therapy. Sci. Robot., 2, eaaq1155 (2017).

43. B. Laycock, M. Nikolić, J.M. Colwell, E. Gauthier, P. Halley, S. Bottle, G. George, Lifetime prediction of biodegradable polymers. Prog. Polym. Sci., 71, 144–189 (2017).

44. P. Calvert, Hydrogels for soft machines. Adv. Mater., 21, 743–756 (2009).

45. J. Li, D. J. Mooney, Designing hydrogels for controlled drug delivery. Nat. Rev. Mater., 1, 16071 (2016).

46. O. Kargiotis, O. Kargiotis, C. Chetty, C. S. Gondi, A. J. Tsung, D. H. Dinh, M. Gujrati, S. S. Lakka, A. P. Kyritsis, J. S. Rao, Adenovirus-mediated transfer of siRNA against MMP-2 mRNA results in impaired invasion and tumor-induced angiogenesis, induces apoptosis in vitro and inhibits tumor growth in vivo in glioblastoma. Oncogene, 27, 4830 (2008).

47. G. J. Hancock, The self-propulsion of microscopic organisms through liquids. Proc. R. Soc. Lond. A. Math. Phys. Sci., 217, 96–121 (1953).

48. A. F. Tabak, S. Yesilyurt, Improved kinematic models for two-link helical micro/nanoswimmers. IEEE Trans. Robot., 30, 14–25 (2014).

49. J. Happel, H. Brenner, Low Reynolds Number Hydrodynamics. NJ: Prentice-Hall (1965).

50. G. I. Taylor, Analysis of the swimming of microscopic organisms. Proc. R. Soc. Lond. A. Math. Phys. Sci., 209, 447–461 (1951).

51. J. Funda, R. H. Taylor, R. P. Paul, On homogeneous transforms, quaternions, and computational efficiency, IEEE Trans. Robot. Autom. 6, 382–388 (1990).

52. I. S. Khalil, A. F. Tabak, A. Hosney, A. Mohamed, A. Klingner, M. Ghoneima, M. Sitti, Sperm shaped magnetıc microrobots: Fabrication using electrospinning, modeling, and characterization. in Proceedings IEEE International Conference on Robotics and Automation, 16 to 21 May 2016, Stockholm, Sweden, pp. 1939–1944.

53. V. E. Baranova, P. F. Baranov, The Helmholtz coils simulating and improved in COMSOL. Proceedings of the Conference on Dynamics of Systems, Mechanisms and Machines (Dynamics) (2014).

54. A. Ghosh, P. Fischer, Controlled propulsion of artificial magnetic nanostructured propellers. Nano Lett., 9, 2243–2245 (2009).

55. B.D. Fairbanks, M.P Schwartz, C.N. Bowman, K.S. Anseth, Photoinitiated polymerization of PEG-diacrylate with lithium phenyl-2,4,6-trimethylbenzoylphosphinate: polymerization rate and cytocompatibility. Biomaterials, 30, 6702–6707 (2009).

56. D. Loessner, C. Meinert, E. Kaemmerer, L.C. Martine, K. Yue, P.A. Levett, T.J. Klein, F.P. Melchels, A. Khademhosseini, D.W. Hutmacher, Functionalization, preparation and use of cell-laden gelatin methacryloyl-based hydrogels as modular tissue culture platforms. Nat. Protoc., 9, 727–746 (2016).

57. M. J. McGirt, D. Mukherjee, K. L. Chaichana, K. D. Than, J. D. Weingart, A. Quinones-Hinojosa, Association of surgically acquired motor and language deficits on overall survival after resection of glioblastoma multiforme. Neurosurgery 65, 463–469 (2009).

58. A. Servant, F. Qiu, M. Mazza, K. Kostarelos, B. J. Nelson, Controlled in vivo swimming of a swarm of bacteria-like microrobotic flagella. Adv.Mater. 27, 2981–2988 (2015).

59. J. Rahmer, C. Stehning, B. Gleich, Spatially selective remote magnetic actuation of identical helical micromachines. Sci. Robot. 2, eaal2845 (2017).

60. B. Mostaghaci, O. Yasa, J. Zhuang, M. Sitti, Bioadhesive bacterial microswimmers for targeted drug delivery in the urinary and gastrointestinal tracts. Adv. Sci. 4, 1700058 (2017).

